# MultiGWAS: An integrative tool for Genome Wide Association Studies (GWAS) in tetraploid organisms

**DOI:** 10.1101/2020.08.16.252791

**Authors:** L. Garreta, I. Cerón-Souza, M.R. Palacio, P.H. Reyes-Herrera

**Affiliations:** Corporación Colombiana de Investigación Agropecuaria (AGROSAVIA), CI Tibaitatá, Kilómetro 14, Vía a Mosquera, 250047, Colombia; Corporación Colombiana de Investigación Agropecuaria (AGROSAVIA), CI El Mira, Kilómetro 38, Vía Tumaco Pasto, Colombia

**Keywords:** GWAS on polyploids, GWASPoly, GAPIT, SNP, SHEsis, software, TASSEL

## Abstract

**Summary:** The Genome-Wide Association Studies (GWAS) are essential to determine the genetic bases of either ecological or economic phenotypic variation across individuals within populations of the model and non-model organisms. For this research question, the GWAS replication testing different parameters and models to validate the results’ reproducibility is common. However, straightforward methodologies that manage both replication and tetraploid data are still missing. To solve this problem, we designed the MultiGWAS, a tool that does GWAS for diploid and tetraploid organisms by executing in parallel four software, two designed for polyploid data (GWASpoly and SHEsis) and two for diploids data (GAPIT and TASSEL). MultiGWAS has several advantages. It runs either in the command line or in a graphical interface; it manages different genotype formats, including VCF. Moreover, it allows control for population structure, relatedness, and several quality control checks on genotype data. Besides, MultiGWAS can test for additive and dominant gene action models, and through a proprietary scoring function, select the best model to report its associations. Finally, it generates several reports that facilitate identifying false associations from both the significant and the best-ranked association SNP among the four software. We tested MultiGWAS with public tetraploid potato data for tuber shape and several simulated data under both additive and dominant models. These tests demonstrated that MultiGWAS is better at detecting reliable associations than using each of the four software individually. Moreover, the parallel analysis of polyploid and diploid software that only offers Multi-GWAS demonstrates its utility in understanding the best genetic model behind the SNP association in tetraploid organisms. Therefore, MultiGWAS probed to be an excellent alternative for wrapping GWAS replication in diploid and tetraploid organisms in a single analysis environment.

## 1 Introduction

The Genome-Wide Association Studies (GWAS) comprise statistical tests that identify which variants through the whole genome of a large number of individuals are associated with a specific trait [Cantor et al., 2010, Begum et al., 2012]. This methodology started with humans and several model plants, such as rice, maize, and *Arabidopsis* [Lauc et al., 2010, Tian et al., 2011, Cao et al., 2011, Korte and Farlow, 2013, Han and Huang, 2013]. Because of the advances in the high-throughput sequencing technology and the decline of the sequencing cost in recent years, there is an increase in the availability of genome sequences of different organisms at a faster rate [Ekblom and Galindo, 2011, Ellegren, 2014]. Thus, the GWAS is becoming the standard tool to understand the genetic bases of either ecologically or economically relevant phenotypic variation for both model and non-model organisms. This increment includes complex species such as polyploids (Figure 1) [Ekblom and Galindo, 2011, Santure and Garant, 2018].

**Figure 1:**
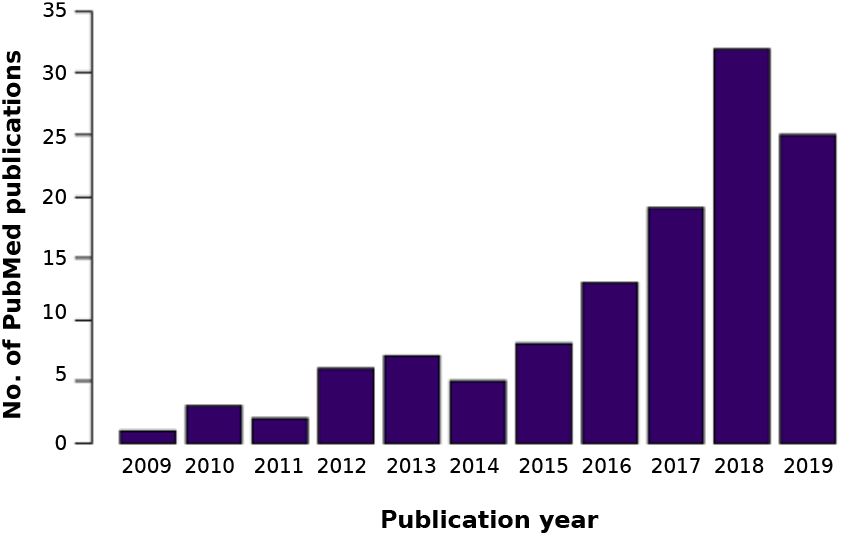
The number of peer-reviewed papers that contains the keywords “GWAS” and “polyploid” in the PubMed database between 2009 and 2019.

The GWAS for polyploid species has fourth related challenges. First, replication among tools is critical to validate GWAS results and capture positive associations [Chanock et al., 2007, De et al., 2014]. Because each tool has its assumptions (i.e., data quality control, false-positive control, and model optimizations, among others), leading to different results such as p-values, significance thresholds, and genomic control inflation factors. Consequently, each tool can be considered an independent environment to replicate a GWAS analysis. For example, the significance threshold for *p-value* changes through four GWAS software (i.e., PLINK, TASSEL, GAPIT, and FaST-LMM) when the sample size varies [Yan et al., 2019]. It means that well-ranked SNPs from one package can be ranked differently in another.

Second, there are very few tools focused on integrating several GWAS software to compare different parameters and conditions across them. As far as we know, there is only two software with this service in mind: iPAT and easyGWAS.The iPAT allows running in a graphic interface three well-known command-line GWAS software such as GAPIT, PLINK, and FarmCPU [Zhang et al., 2018]. However, the output from each package is separated. On the other hand, the easyGWAS allows running a GWAS analysis on the web using different algorithms and combining several GWAS results. This analysis runs independently of both the computer capacity and the operating system. Nevertheless, it needs either several datasets to obtain the different GWAS results to make replicates or GWAS results already computed. In either case, the results from different algorithms are also separated [Grimm et al., 2017]. Thus, although both software iPAT and easyGWAS integrate with different programs or algorithms, an output that allows to compare similitude and differences in the association is missing.

Third, although there are different GWAS software available to repeat the analysis under different conditions [Gumpinger et al., 2018], most of them are designed exclusively for the diploid data matrix [Bourke et al., 2018]. Therefore, it is often necessary to “diploidizing” the polyploid genomic data to replicate the analysis. This process could withdraw how allele dosage affects the phenotype expression in polyploid species [Ferrão et al., 2018]. However, some genome sections of autopolyploid species did not duplicate, leading to loci’s disomic inheritance [Ohno, 1970, Lynch and Conery, 2000, Dufresne et al., 2014]. Moreover, the inheritance mechanism of most of the polyploid species is still unknown. Therefore, software that accounts for both polyploid and diploid data facilitates analyzing both inheritance types in polyploids.

Finally, for polyploid species, any tool that integrates and compares different gene action among software is key to understanding how redundancy or complex interaction among alleles affects the phenotype expression and the evolution of new phenotypes [Bourke et al., 2018, Rosyara et al., 2016, Ferrão et al., 2018].

This study developed the MultiGWAS tool that performs GWAS analyses for both diploid and tetraploid species using four software in parallel to overcome these challenges. The tool includes GWASpoly [Rosyara et al., 2016] and SHEsis [Shen et al., 2016] that accept polyploid genomic data. Also it includes and GAPIT [Tang et al., 2016] and TASSEL [Bradbury et al., 2007], designed for GWAS in plants, but that in the case of tetraploid data, their use require “diploidizing” genomic matrix. This wrapping tool deals with different input file formats that come from several polyploid genotypes calling software, including VCF. Besides, MultiGWAS manages data preprocessing, searches for associations by running four GWAS software in parallel, and creates a score to choose between gene action models in GWASPoly and TASSEL. This study describes the utilities of MultiGWAS and its evaluation through simulation studies and one public GWAS data set, demonstrating its advantages.

## 2 Method

The MultiGWAS tool has three main steps: the adjustment, the multi analysis, and the integration (Figure 2). In the adjustment step, MultiGWAS processes the configuration file. Then it cleans and filters the genotype and phenotype datasets, and in the case of tetraploids, MultiGWAS “diploidize” the genomic data. Next, during the multi-analysis, each GWAS tool runs in parallel. Subsequently, in the integration step, the MultiGWAS tool scans the output data files from the four packages (i.e., GWASPoly, SHEsis, GAPIT, and TASSEL); post-processes the data; and finally, it generates a summary of all results that contain: associations tables, Venn diagrams showing associated SNPs shared among tools, SNPs in linkage disequilibrium (LD), Manhattan and quantile-quantile (QQ) plots, chord diagrams showing the position in the chromosome of the associated SNPs and finally SNP profiles (see Section 2.3.3).

**Figure 2:**
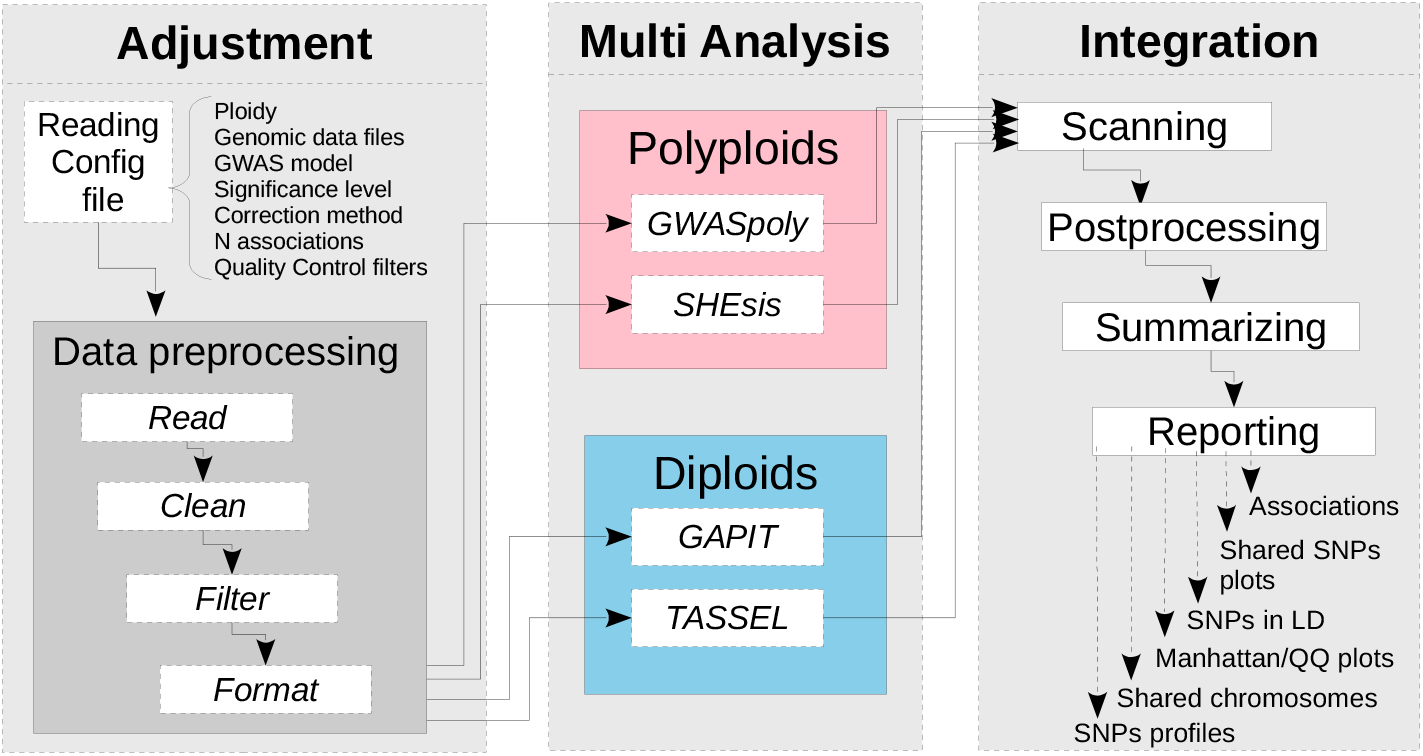
MultiGWAS flowchart has three steps: adjustment, multi analysis, and integration. In the first step, after the input data management upload, MultiGWAS read the configuration file and preprocessing the input data (genotype and phenotype dataset). The second step is the GWAS analysis, where MultiGWAS configure and run the four packages in parallel. Finally, in the third step, MultiGWAS summarize the results and generate a report using different tabular and graphical visualizations.

### 2.1 Adjustment stage

MultiGWAS takes as input a configuration file where the user specifies the genomics data and the parameters for the four tools. Once the configuration file is read and processed, the genomic data files (genotype and phenotype) are then cleaned, filtered, and checked for data quality. The output of this stage corresponds to the inputs for the four programs at the Multi Analysis stage.

#### 2.1.1 Reading configuration file

The configuration file includes the following settings that we briefly describe:

##### Ploidy

Currently, MultiGWAS supports diploid and tetraploid genotypes were 2 for diploids and 4 for tetraploids.

##### Genomic input files

MultiGWAS mainly uses two input files for the genotype and the phenotype files, and depending on the genotype format (see below) could be needed a map file with marker information (chromosome name, genomic position, reference allele, and alternate allele)

For genotypes aligned with a reference genome, specify the N chromosomes/-contigs displayed in the plots. Chromosomes are sorted in decreasing order by size. Chromosome/contig size is approximated to the highest variant position. And, when chromosome/contig names are numerical or are too large, they are changed with the string prefix “contig” and a sequential number from 1 to N.

MultiGWAS allows genotype data in five different formats: “gwaspoly” [Rosyara et al., 2016], “vcf” [Team, 2015][Parra-Salazar et al., 2020], “matrix”, “fitpoly” [Voorrips and Gort, 2018], and “updog” [Gerard and Ferrão, 2020] for-mats. The former two already include marker information, but the last three formats do not, and they need the additional map file. VCF files are transformed into GWASpoly format using NGSEP 4.0.2 [Tello et al., 2019]. A detailed information of these files and formats are available in the Github of the tool (https://github.com/agrosavia-bioinfo/MultiGWAS).

##### Test model

One of the main factors in detecting real trait-marker associations depends on the gene action models. A unique feature offered by Multi-GWAS is to test the different gene action models supported by the tools (see section 2.2). The additive model is the default model supported by all the tools. However, GWASpoly supports eight, TASSEL three, GAPIT two, and SHEsis only supports one. To integrate the different models in one wrapping tool, MultiGWAS offers three testing options: “additive” (supported by all the tools), “dominant” (supported by all tools except SHEsis), and “all” (for testing all effects supported by the tools, including both additive and dominant effects). In any of the three tests, MultiGWAS reports its top N associations with low *p-value* (With N defined by the user, see below). Taking these associations is straightforward for the former two tests, but not for the last one, as tools report different associations for each gene action model. For this last test, we have created a method that automatically selects the “best” gene action model described in Section 2.3.1.

##### GWAS model

MultiGWAS works with quantitative phenotypes and runs two types of GWAS, either with control for population structure and relatedness between samples or without any control. The first is known in the literature as the Q+K or *full model*, where Q refers to population structure and K to relatedness; and the second is known as the *naive model* [Sharma et al., 2018].

Both models are linear regression approaches, and GWAS tools used by MultiGWAS implements some variations of those models. The *naive* is modeled with Generalized Linear Models (GLMs, Phenotype + Genotype), and the *full* is modeled with Mixed Linear Models (MLMs, Phenotype + Genotype + Structure + Kinship). The default model used by MultiGWAS is the *full model* (Q+K) [Yu et al., 2006], following the equation:

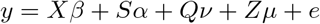

In this equation, the *y* is the vector of observed phenotypes. Moreover, the *β* is a vector of fixed effects other than SNP or population group effects, the *α* is a vector of SNP effects (Quantitative Trait Nucleotides), the *ν* is a vector of population effects, the *µ* is a vector of polygene background effects, and the *e* is a vector of residual effects. Besides, *Q*, modeled as a fixed effect, refers to the incidence matrix for subpopulation covariates relating *y* to *ν*, and *X, S* and *Z* are incidence matrices of 1s and 0s relating *y* to *β, α*, and *µ*, respectively.

##### Genome-wide significance

GWAS searches SNPs associated with the phenotype in a statistically significant manner. A threshold or significance level *α* is specified and compared with the *p-value* derived for each association score. Standard significance levels are 0.01 or 0.05 [Gumpinger et al., 2018, Rosyara et al., 2016], and MultiGWAS uses an *α* of 0.05 for the four GWAS packages. However, in GWASpoly and TASSEL, which calculates the SNP effect for each genotypic class using different gene action models (see “Multi analysis stage”), the threshold is adjusted according to these two packages. Therefore, the number of tested markers may be different in each model (see below), impacting the *p-value* thresholds.

##### Multiple testing correction

Due to the massive number of statistical tests performed by GWAS, it is necessary to perform a correction method for multiple hypothesis testing and adjusting the *p-value* threshold accordingly. Two standard methods for multiple hypothesis testing are the false discovery rate (FDR) and the Bonferroni correction. The latter is the default method used by MultiGWAS, which is one of the most rigorous methods. However, instead of adjusting the *p-values*, MultiGWAS adjust the threshold below which a *p-value* is considered significant. That is *α/m*, where *α* is the significance level, and *m* is the number of tested markers from the genotype matrix.

##### Number of reported associations

The use of stringent significance levels could discard many *p-value* associations closer to significant threshold, generating a high number of false negatives [Thompson et al., 2011, Kaler and Purcell, 2019]. To avoid this problem, MultiGWAS provides the option to specify the number of best-ranked associations (lower *p-values*), adding the corresponding *p-value* to each association found. In this way, it is possible to enlarge the number of results and their replicability across the different programs. Nevertheless, the report displays each association with its corresponding *p-value*.

##### Quality control filters

A control step is necessary to check the input data for the genotype or phenotype errors or low quality, leading to spurious GWAS results. MultiGWAS provides the option to select and define thresholds for the following filters that control the data quality: Minor Allele Frequency (MAF), individual missing rate (MIND), SNP missing rate (GENO), and Hardy-Weinberg threshold (HWE). All of these filters are built-in implementations of multiGWAS, except the HWE for tetraploids:

- **MAF of *x:*** filters out SNPs with minor allele frequency below *x* (default 0.01);
- **MIND of *x:*** filters out all individuals with missing genotypes exceeding *x*\*100% (default 0.1);
- **GENO of *x:*** filters out SNPs with missing values exceeding *x*\*100% (default 0.1);
- **HWE of *x:*** filters out SNPs with a *p-value* below the *x* threshold in the Hardy-Weinberg equilibrium exact test. In the case of tetraploid genotypes this calculation is taken from SHEsis [Shen et al., 2016].

##### GWAS tools

List of the four GWAS software names to run and integrate into MultiGWAS analysis: GWASpoly and SHEsis (designed for polyploid data), and GAPIT and TASSEL (designed for diploid data).

##### Linkage Disequilibrium threshold (*R*^2^)

User-defined squared correlation threshold (*R*2) above which a pair of SNPs is considered to be in linkage disequilibrium (see Section 2.3.3 for details).

#### 2.1.2 Data preprocessing

Once the configuration file is processed, the genomic data is read and cleaned by selecting individuals present in both genotype and phenotype. Then, Multi-GWAS removes individuals and SNPs with low quality following the previous selected quality-control filters and their thresholds,

At this point, the format “ACGT” suitable for the polyploid software GWAS-poly and SHEsis, is “diploidized” for GAPIT and TASSEL. The homozygous tetraploid genotypes are converted to diploid thus: AAAA→AA, CCCC→CC, GGGG→GG, TTTT→TT. Moreover, for tetraploid heterozygous genotypes, the conversion depends on the reference and alternate alleles calculated for each position (e.g., AAAT →AT,…, CCCG→CG).

After this process, MultiGWAS converts the genomic data, genotype, and phenotype datasets to the specific formats required for each of the four GWAS packages.

### 2.2 Multi analysis stage

As described in Section 2.1.1, MultiGWAS can run two types of GWAS: *naive* without any genotype data control and *full* with control for population structure and relatedness. GWASpoly, GAPIT, and TASSEL support both models. However, SHEsis, supports only the *naive* model. To control population structure and relatedness in the *full* model, MultiGWAS uses built-in algorithms to calculate both principal components as covariates and kinship among pairs of individuals. A more detailed description of each of the GWAS tools is below.

#### 2.2.1 GWASpoly

GWASpoly [Rosyara et al., 2016] is an R package designed for GWAS in polyploid species used in several studies in plants [Berdugo-Cely et al., 2017, Ferrão et al., 2018, Sharma et al., 2018, Yuan et al., 2019]. GWASpoly uses a Q+K linear mixed model with biallelic SNPs that account for population structure and relatedness. Also, to calculate the SNP effect for each genotypic class, GWASpoly provides eight gene action models: general, additive, simplex dominant alternative, simplex dominant reference, duplex dominant alternative, duplex dominant, diplo-general, and diplo-additive. Consequently, the number of statistical tests performed can be different in each action model, and thus thresholds below which the *p-values* are considered significant.

MultiGWAS is using GWASpoly version 1.3 with all gene action models available to find associations. The MultiGWAS reports the top *N* best-ranked (the SNPs with lowest *p-values*) that the user specified in the *N* input configuration file. The *full* model used by GWASpoly includes the population structure and relatedness, which are estimated using the first five principal components and the kinship matrix, respectively, both calculated with the GWASpoly built-in algorithms.

#### 2.2.2 SHEsis

SHEsis [Shen et al., 2016] is a program based on a linear regression model that includes single-locus association analysis for polyploids, among other analyses. However, it has been used mainly by animals and humans, both diploids [Qiao et al., 2015, Meng et al., 2019].

MultiGWAS uses version 1.0, which does not take account of population structure or relatedness. However, MultiGWAS externally estimates relatedness for SHEsis by excluding individuals with cryptic first-degree relatedness using the kinship matrix calculated by GWASpoly built-in algorithm.

#### 2.2.3 GAPIT

GAPIT is an R-based program designed for plants. This tool implements the classical MLM for the *full* model correcting by population structure and relatedness. Also, it uses the GLM approach for the *naive* model without any correction [Tang et al., 2016].

GAPIT offers two models of gene action: additive and dominant. For both models, the genotype must be in numerical format. For the additive model, the genotype is implicitly transformed by GAPIT, using 0 for homozygous genotypes with recessive allele combinations, 2 for homozygous genotypes with dominant allele combinations, and 1 for heterozygous genotypes. For the dominant model, MultiGWAS transforms the genotype, using 0 for the two types of homozygous genotypes and 1 for heterozygous genotypes, as indicated by the authors [Tang et al., 2016]. MultiGWAS uses the latest version 3, which also implements several state-of-the-art methods developed for statistical genomics [Wang and Zhang, 2020].

#### 2.2.4 TASSEL

TASSEL is another standard GWAS program developed initially for maize but currently used in several species [Álvarez et al., 2017, Zhang et al., 2018]. TASSEL is a java package that runs either using a graphic user interface developed in JAVA or a command-line interface through a Perl pipeline. In MultiGWAS is implemented the Perl pipeline.

For the association analysis, TASSEL includes the general linear model (GLM) for a naive analysis. Moreover, it uses the mixed linear model (MLM) for a full analysis controlling for population structure, a principal component analysis, and controlling relatedness using a kinship matrix with a centered IBS method with TASSEL built-in algorithms. Moreover, as GWASPoly, TASSEL provides three-gene action models to calculate each genotypic class’s SNP effect: general, additive, and dominant. Hence, the significance threshold depends on each action model.

### 2.3 Integration stage

The outputs resulting from the four GWAS packages are post-processed to identify SNP with either significative *p-values* association or best-ranked association (i.e., with *p-values* close to a significance threshold).

#### 2.3.1 Selection of best gene action model

MultiGWAS offers three testing options: “additive”, “dominant”, and “all”. Taking the best associations from “additive” and “dominant” tests is straight-forward. However, for the option “all”, MultiGWAS has a method to select within each tool the “best” gene action model and takes the top associations.

The method works by scoring each gene action model using three criteria: inflation factor (I), shared SNPs (R) and significant SNPs (S), using the following equation:

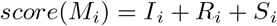

where *score*(*M*_*i*_) is the score for the gene action model *M*_*i*_, with *i* from 1..*k*, for a GWAS package with *k* gene action models. *I*_*i*_ is the score for the inflation factor defined as *I*_*i*_ = 1 − |1 − λ(*M*_*i*_)|, where λ(*M*_*i*_) is the inflation factor for the *M*_*i*_ model. *R*_*i*_ is the score of the shared SNPs defined as 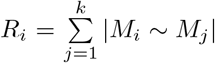 where |*M*_*i*_ ∼ *M*_*j*_|, is the number of SNPs shared between *M*_*i*_ and *M*_*j*_ models, normalized by the maximum number of SNPs shared between all models. And, *S*_*i*_ is the number of significant SNPs of model *M*_*i*_ normalized by the total number SNPs shared among all models.

The score is high when an *M*_*i*_ model has an inflation factor λ close to 1, identifies a high number of shared SNPs, and contains one or more significant SNPs. Conversely, the score is low when the *M*_*i*_ model has an inflation factor λ either low (close to 0) or high (λ > 2), which identifies a small number of shared SNPs, and contains 0 or few significant SNPs. In any other case, the score results from the balance among the inflation factor, the number of shared SNPs, and the number of significant SNPs.

#### 2.3.2 Selection of significant and best-ranked associations

MultiGWAS reports two groups of associations from the four GWAS packages: the statistically significant associations with *p-values* below a threshold of significance, and the best-ranked associations with the lowest *p-values*, but not reaching the limit to be statistically significant. However, they are representing interesting associations for further analysis (possible false negatives).

#### 2.3.3 Integration of results

All four GWAS packages adopted by MultiGWAS use linear regression approaches. However, they often produce different association results for the same input. Computed *p-values* for the same set of SNPs are different between pack-ages. Therefore, SNPs with significant *p-values* for one package maybe not significant for the others. Alternatively, well-ranked SNPs in one package may be ranked differently in another.

MultiGWAS integrates the results of the four tools generating six types of outputs that combine graphics and tables to compare, select, and interpret the set of possible SNPs associated with a trait of interest (Fig 3). The unified output is one HTML document that contains the tables and figures to cover all user’s needs to present results and includes:

**Figure 3:**
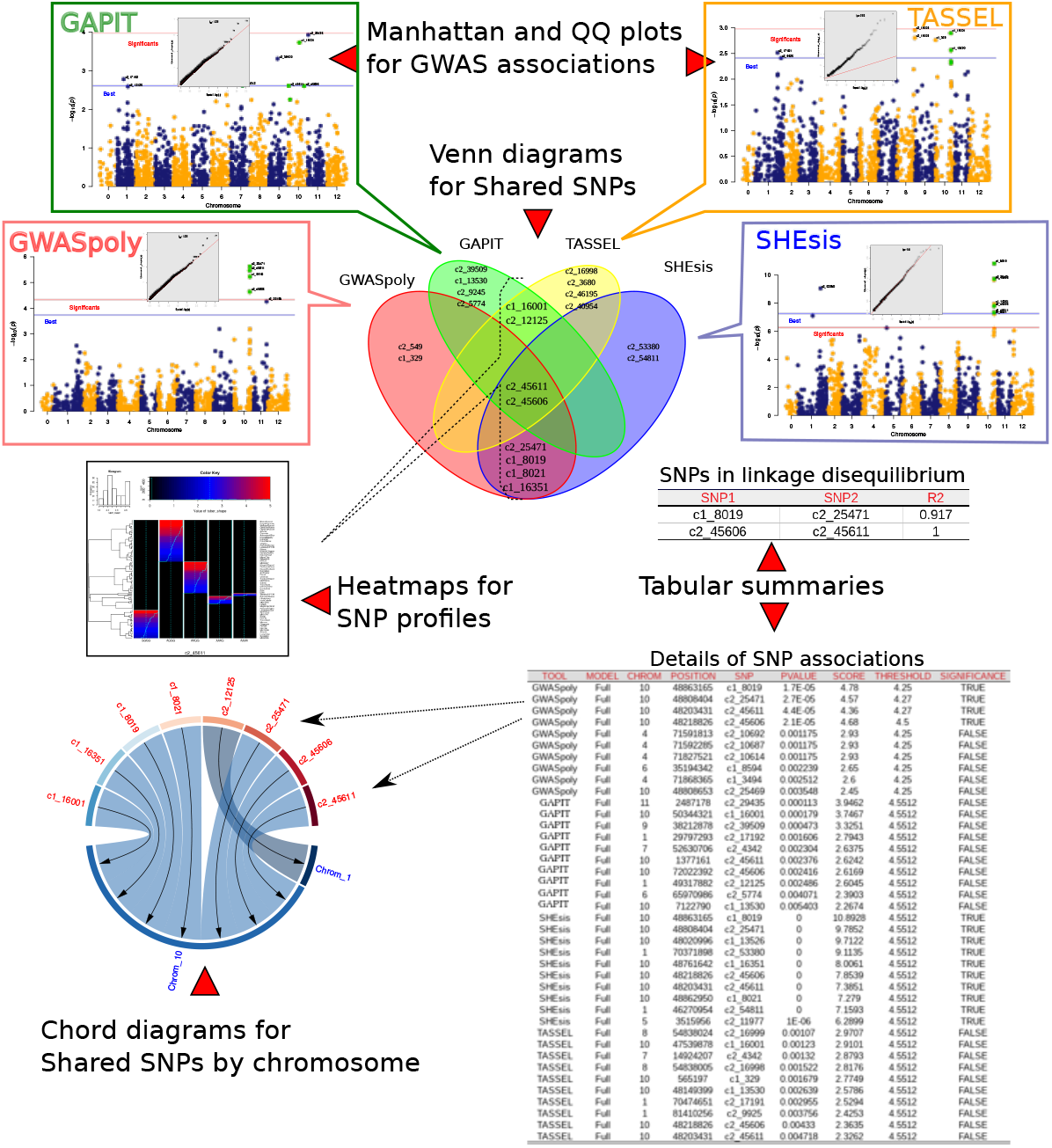
Reports presented by MultiGWAS. Results for GWASpoly, GAPIT, TASSEL, and SHEsis. For each tool we present, first, a QQ plot that assesses the resultant *p-values*, and second, a Manhattan plot with two lines, blue and red, representing the lower limit for the best ranked and significant SNPs, respectively. Also, we present Venn diagrams that visualize the reproducibility of results; SNPs profiles that visualize SNPs by a two-dimensional representation; chord diagrams that show how the strongest associations are limited to a few chromosomes. Furthermore, we present tabular summaries with details of SNPs in linkage disequilibrium and significant and best-ranked associations.

##### QQ plots for GWAS associations

The QQ plot shows how well most SNPs fit the null hypothesis of no association with the phenotype. Both distributions should coincide, and most SNPs should lie on the red diagonal line. Deviations for many SNPs may reflect inflated *p-values* due to population structure or cryptic relatedness. Nevertheless, few SNPs deviate from the diagonal for a truly polygenetic trait [Power et al., 2016]. MultiGWAS adds the top of each QQ plot the corresponding inflation factor λ to assess the test statistic inflation degree.

##### Manhattan plot for GWAS associations

MultiGWAS uses classical Manhattan plots to visualize each package’s results. In both plots, the points are the SNPs and their *p-values* are transformed into scores like −*log*_10_(*p*-*values*) (see Figure 3). The Manhattan plot shows the strength of association of the SNPs (y-axis) distributed at their genomic location (x-axis), so the higher the score, the stronger the association. MultiGWAS adds distinctive marks to the plot; significant SNPs are above a red line, best-ranked SNPs are above a blue line, and SNPs shared between packages are colored green.

##### Tables and Venn diagrams for single and shared SNPs

MultiGWAS provides tabular and graphic views to report the best-ranked and significant SNPs identified by the four GWAS packages in an integrative way (Figure 3). Both *p-values* and significance levels have been scaled as −*log*_10_(*p*-*values*) to give high scores to the best statistically evaluated SNPs.

First, best-ranked SNPs correspond to the top-scored *N* SNPs, whether they were assessed significant or not by its package, and with *N* defined by the user in the configuration file. These SNPs appear in both an SNPs table and in a Venn diagram. The table lists them by package and sorts by decreasing score, whereas the Venn diagram emphasizes if they were best-ranked either in a single package or in several at once (shared).Second, the significant SNPs correspond to the ones valued statistically significant by each package. They also appear in a Venn diagram and the SNPs table, marked with significance TRUE (T).

##### Views of SNPs in linkage disequilibrium (LD)

MultiGWAS reports a Venn diagram and a table (Figures 7.a and 7.b, respectively) for pairs of SNPs with squared correlation equal to or greater than the threshold *R*2, where *R*2 is defined by the user in the configuration file (see 2.1.1). MultiGWAS joins the N best associations found for each GWAS packages (SNPs with the lowest *p-value*), calculates for each pair of SNPs the *R*2 using the R ldsep library for LD in polyploids [Gerard, 2021]. Finally, it summarizes the results in a table with pairs of SNPs per row along with their calculated *R*^2^.

**Figure 7:**
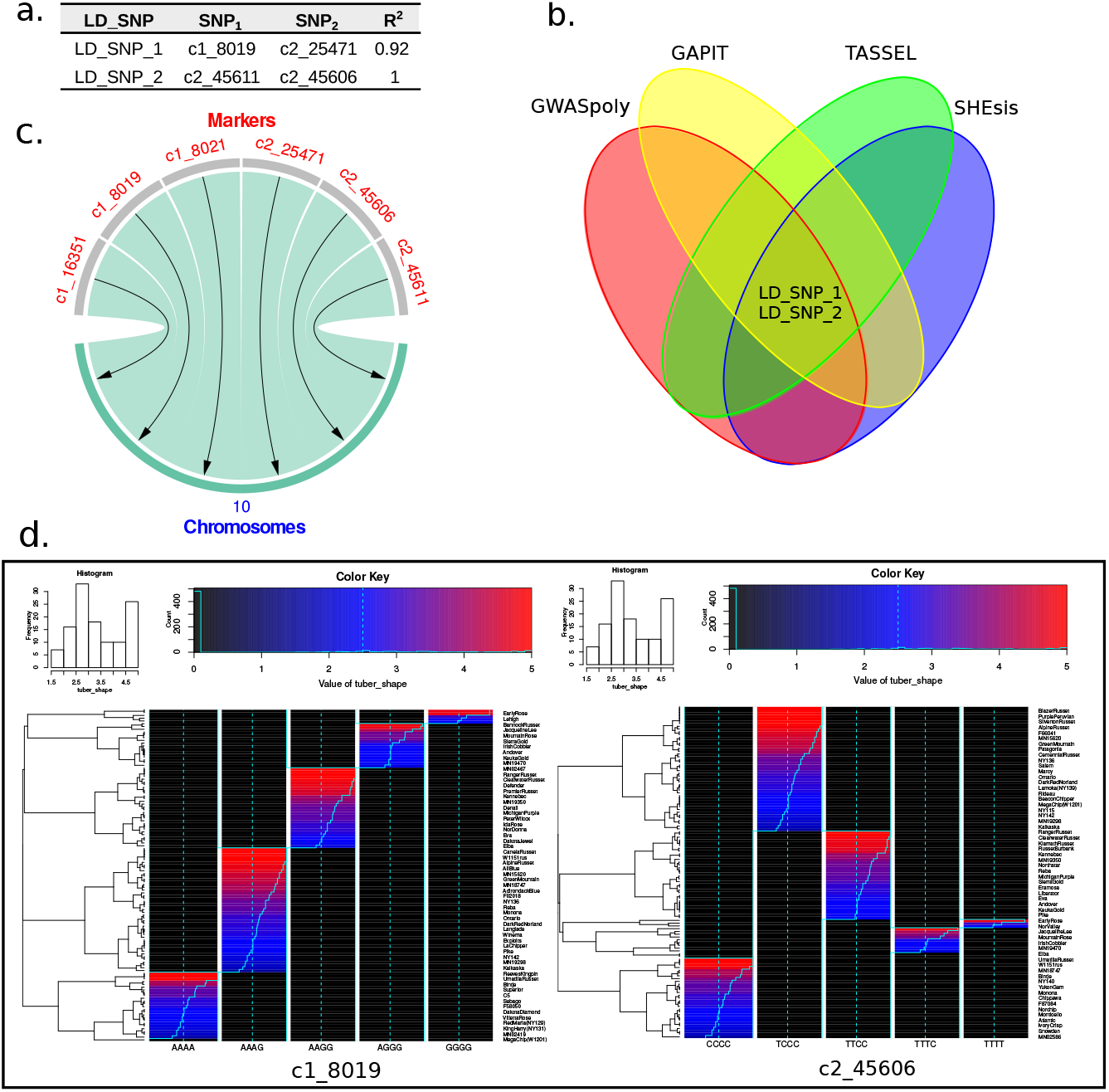
SNPs in LD, SNPs by chromosome, and SNPs structure. MultiGWAS brings different views to figure out SNP relationships and structure. **a**. Table of pairs of SNPs in LD with columns LD_SNP for an ID given by MultiGWAS to the pair of SNPs; SNP1 and SNP2 for the names of SNPs; and *R*^2^ for the squared correlation value between SNPs. **b**. Venn diagram of pairswise SNPs in LD. LD_SNPs in the center show that the four packages simultaneously detected at least one SNP from the pairwise SNPs shown in the table. **c**. Chord diagram showing that best-ranked SNPs located in chromosome 10. **d**. SNP profiles showing the structure of one of the pairwise SNPs in LD.

Pairs of SNPs in LD are assigned a new ID (LD_SNP) and reported in a Venn diagram highlighting the shared SNPs in LD detected between the GWAS software. This view allows for quick identification of related SNPs with different names instead of a plain table, as most GWAS packages report their results.

##### Heatmaps for the structure of shared SNPs

For each SNP identified more than once, MultiGWAS provides its SNP profile. It is a two-dimensional heatmap representing the SNP that visualizes each trait by individuals and genotypes as rows and columns, respectively.

Within the figure, at the left, the individuals are grouped in a dendrogram by their genotype. At the right, there is the name or ID of each individual. At the bottom, the genotypes are ordered from left to right, starting from the major to the minor allele (i.e., AAAA, AAAB, AABB, ABBB, BBBB). At the top, there is a description of the trait based on a histogram of frequency (top left) and an assigned color for each numerical phenotype value using a color scale (top right). Thus, each individual appears as a colored line by its phenotype value on its genotype column. For each column, there is a solid cyan line with each column’s mean and a broken cyan line that indicates how far the cell deviates from the mean (Figure 3).

Because each multiGWAS report shows one specific trait at a time, the histogram and color key will remain the same for all the best-ranked SNPs.

##### Chord diagrams for SNPs by chromosome

The chord diagrams visualize the location across the genome of the best-ranked associated SNPs shared among the four packages and described in the tables. This visualization complements the Manhattan plots from each GWAS package (Figure 3).

## 3 Availability and Implementation

MultiGWAS is a wrapping tool developed in R (R>=3.6). However, it is not an R package or run in the R interface. Instead, it runs on Linux environments because it integrates four external GWAS software implemented in different languages. GWASpoly and GAPIT are R packages; SHEsis is a binary program developed in C++, and TASSEL is a java package that runs through a pipeline implemented in Perl. Consequently, users can run MultiGWAS either by a command-line interface (an R script) or a graphic user interface (a Java application). For detailed instructions and usage examples, refer to https://github.com/agrosavia-bioinfo/MultiGWAS#running-the-examples.

### 3.1 Input parameters

MutiGWAS uses a single configuration text file with the values for the main parameters that drive the analysis. If users prefer a text file, it must have the parameter names and values separated by a colon, filenames without blank spaces, TRUE or FALSE values to indicate if filters are applied, and NULL value to indicate that there is no value for the parameter. This file must have the structure showed in Figure 4. In contrast, if users prefer the GUI application, they can create the configuration file using the GUI described in section 3.2.2.

**Figure 4:**
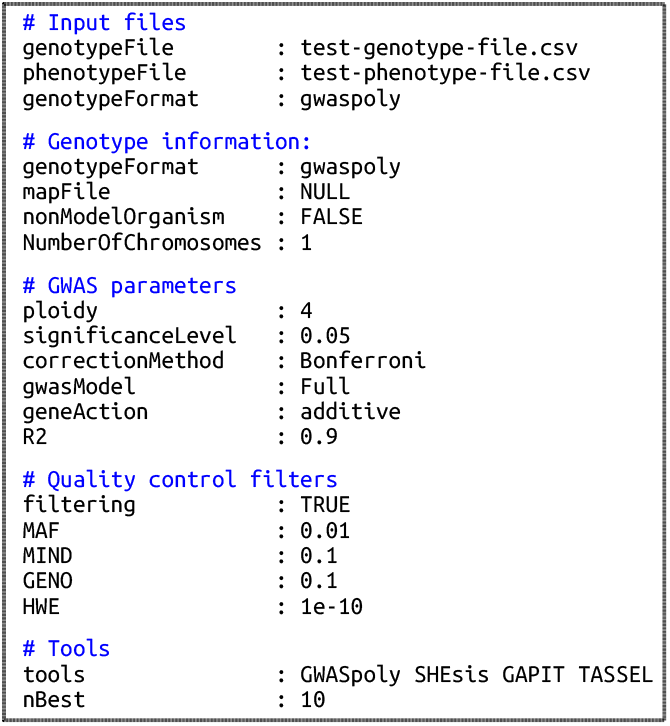
Example of the configuration text file for running MultiGWAS. The input parameters have five sections: (1) Input files for genotype and phenotype file paths. (2) Genotype information for additional information of the genotype. (3) GWAS parameters for setting the main parameter driving the GWAS analysis. (4) Quality control filters to enable/disable and set values for quality control filters on the data. (5) Tools for setting the software to include in the analysis and the number of associations to be reported. The details of each parameter are in Section 2.1.1.

The input files (genotype/phenotype/map) do not need to be in the working directory, but if this is the case, MultiGWAS needs the absolute path.

### 3.2 Installing and using MultiGWAS

MultiGWAS offers different installations: from scratch, precompiled versions, a virtual machine, and a docker image. Specific instructions for the different installation types, including a ready-to-use Linux virtual machine (VM) for running MultiGWAS on other platforms (Windows, OS X), are available in the Github of the tool (https://github.com/agrosavia-bioinfo/MultiGWAS).

#### 3.2.1 Using the command line interface

The execution of the CLI tool is simple. In a Linux console, move to the folder where is the configuration file, and type the executable tool’s name, followed by the filename of the configuration file, like this:

`multiGWAS Test01.config`

Then, the tool starts the execution, showing information on the process in the console window. When it finishes, the results are in a new subfolder called *“out-Test01*, containing a subfolder for each trait in the phenotype file. The results in each trait subfolder include a complete HTML report containing the different views described in the methods section, the source graphics and tables supporting the report, and the preprocessed tables from the results generated by the four GWAS packages used by MultiGWAS.

#### 3.2.2 Using the graphical user interface

The interface allows users to save, load, or specify the different input parameters for MultiGWAS in a friendly way (Figure 5). The input parameters correspond to the settings included in the configuration file described in subsection 2.1.1. It executes by calling the following command from a Linux console:

**Figure 5:**
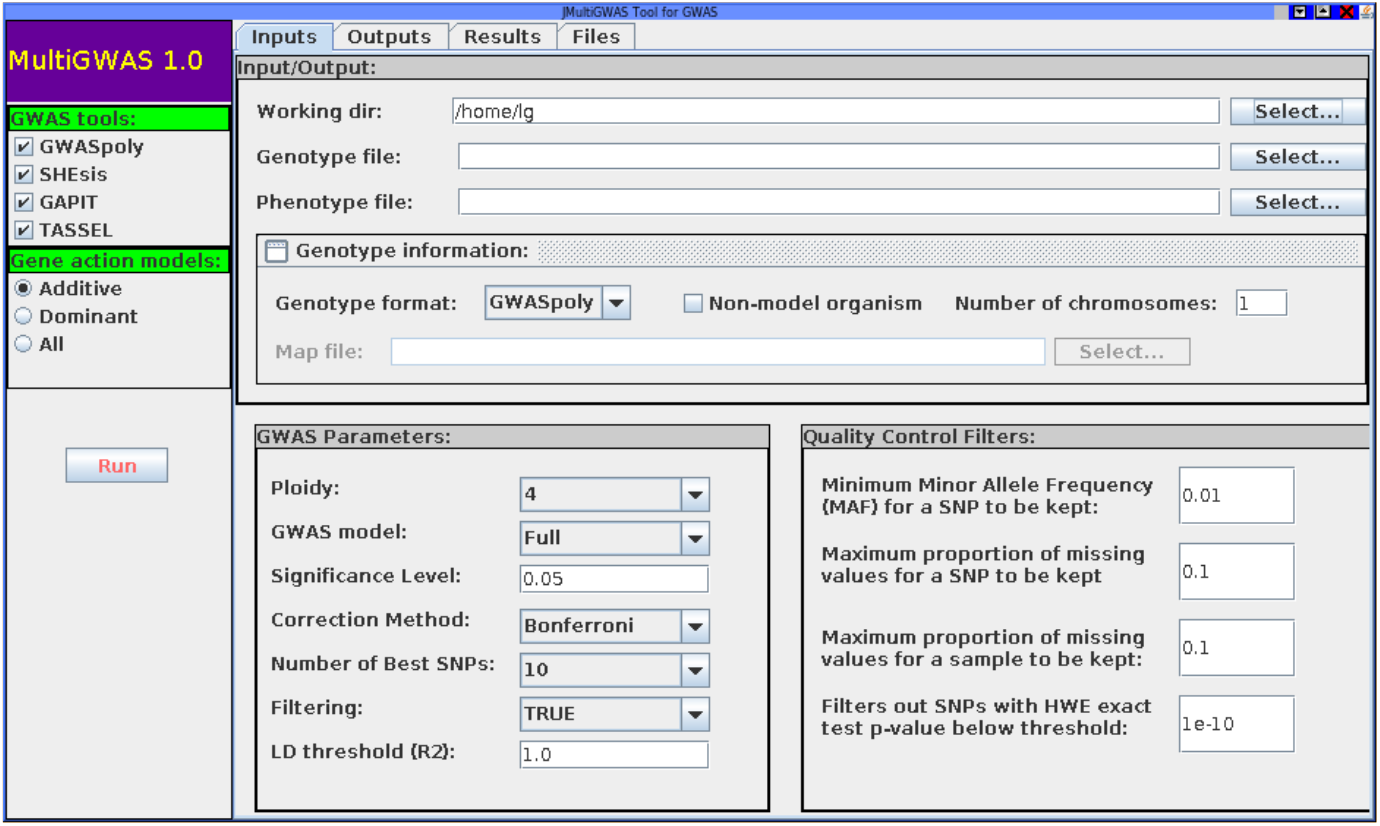
Main view of the MultiGWAS graphical user interface. The interface has a toolbar at the left side and four tabs at the top. In the toolbar, users can select the GWAS tools (GWASpoly, SHEsis, TASSEL, GAPIT). In the Input tab, at top, users can specify the working directory where outputs will be saved, along with genotype, phenotype and additional information of the genotype. And at bottom, users can set the GWAS parameters and quality control filters. The Output tab shows the execution of each process. In the Results tab, users can browse the HTML report of the current analysis generated by the tool. Finally, in the Files tab, users can browse the source files of each software and access the produced data across the analysis. The details of each parameter are described in Section 2.1.1.

`jmultiGWAS`

## 4 Testing MultiGWAS

### 4.1 Testing MultiGWAS in real data

We tested MultiGWAS in real data using an open dataset of a diversity panel of phenotype and genotype information for tetraploid potato. This data is part of the USDA-NIFA SOLCAP Project [Hirsch et al., 2013]. We limited the experiment only for the tuber shape trait, testing both the full and the naive GWAS model.

### 4.2 Testing MultiGWAS in simulated data

We created two different simulated genotyping-phenotyping datasets as an experiment to determine the advantages of running a wrapping tool as MultiGWAS compared with an individual analysis of each of the four GWAS software that integrate MultiGWAS (i.e., GWASpoly, SHEsis, GAPIT, and TASSEL). The first simulated data set had an additive inheritance model, and the second one had a dominant inheritance model.

In both simulations, we used as a founder population a subset of 400 SNP and 150 individuals from tetraploid potato data described by [Enciso-Rodriguez et al., 2018]. To create both simulations of phenotypes, we sampled either additive or dominant effects from a gamma distribution *Γ* (shape = 0.2 and scale = 5) and specified ten SNPs as causal SNPs along with their effects under the Phyton3 SeqBreed software [Pérez-Enciso et al., 2020], inspired in the pSBVB software created to generate polyploid data [Zingaretti et al., 2019].

Both simulated data sets were analyzed in MultiGWAS using the following parameters: Gene action, either additive or dominant, false filtering, Bonferroni correction method, and Naive GWAS model using the founder genotype and either the additive or dominant phenotype. After MultiGWAS analysis, we summarized the top SNPs (i.e., the N best-ranked SNPs found by the tool) and significant SNPs found by each GWAS tool. Then, we calculated two metrics: true-positive rates (TPR) and true-negative rates (TNR) expressed in the following equations:

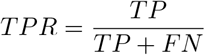

Where TP is the number of SNPs correctly identified as causal SNPs, and FN is the number of SNPs incorrectly identified as non-causal SNPs.

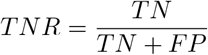

Where TN is the number of SNPs correctly identified as non-causal SNPs, and FP is the number of SNPs incorrectly identified as causal SNPs.

## 5 Results

### 5.1 MultiGWAS performance in real data

We run MultiGWAS for the tuber shape of the tetraploid potato dataset using a full GWAS model controlling the population structure and relatedness [Hirsch et al., 2013].

The full GWAS analysis found several associated SNPs (table of Figure 6.a). From them, three SNPs named as c2_25471, c2_45606, and c2_45611, were detected from top SNPs across the four GWAS packages (central intersection in the Figure 6.a). Two SNPs, named as c1_8019 and c2_25471, were identified as significant by the polyploid packages GWASpoly and SHEsis (Figure 6.b). Previous association studies also reported these SNP where the SNP c1_8019 is associated with potato tuber shape and eye depth traits [Rosyara et al., 2016, Sharma et al., 2018], while the SNPs c2_45606 and c2_45611 are associated to eye depth [Totsky et al., 2020].

**Figure 6:**
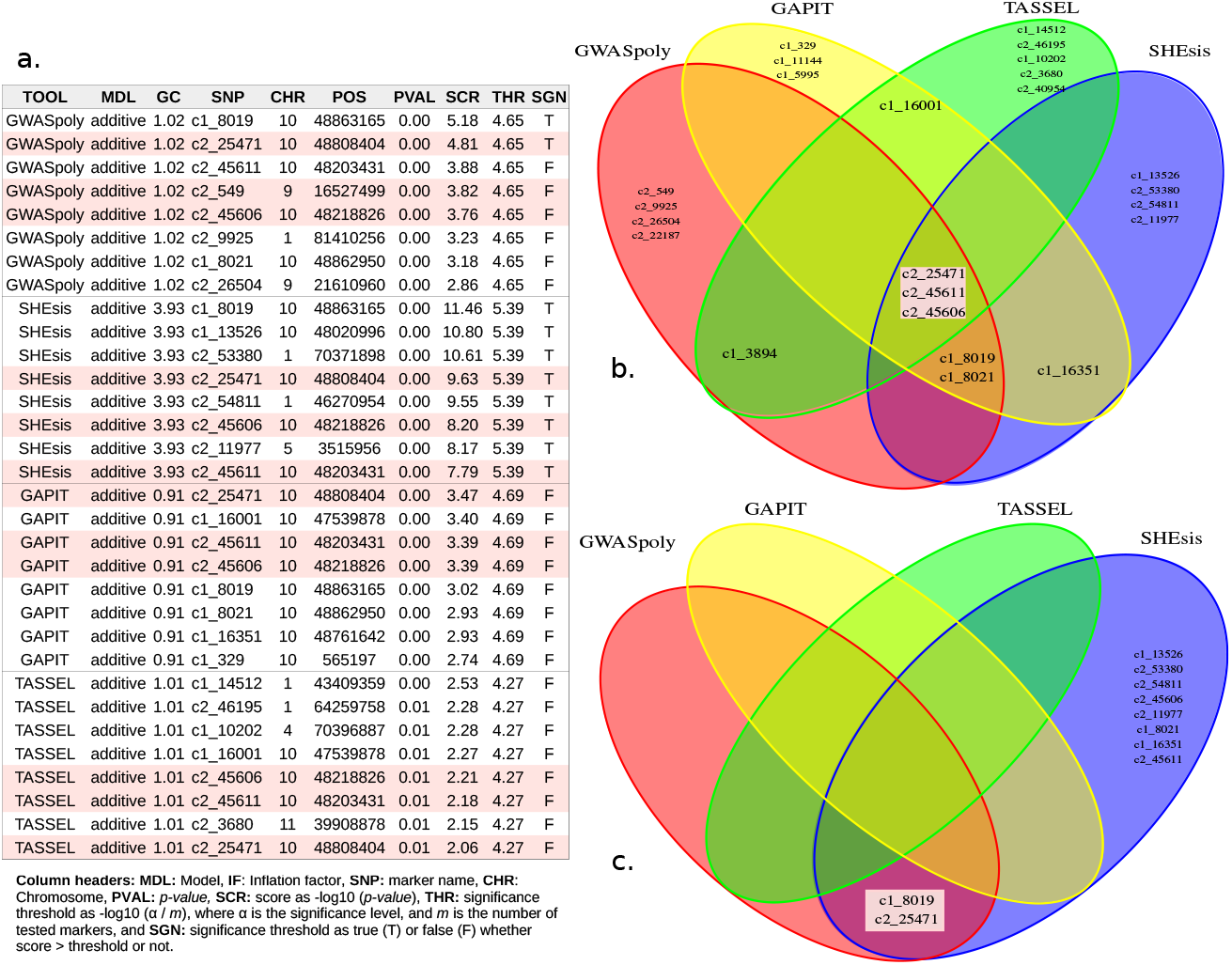
Shared SNPs Views. Tabular and graphical views of SNP associations identified by one or more GWAS packages (shared SNPs). SNPs identified by all packages are marker with red background in all figures. **a**. Table with details of the N=8 best-ranked SNPs from each GWAS package. Each row corresponds to a single SNP. **b**. Venn diagram of the best-ranked SNPs. SNPs identified by all packages are in the central intersection. Other shared SNPs are in both upper central and lower central intersections. **c**. Venn diagram of the significant SNPs (score threshold).

MultiGWAS strengthened the replicability of these associated SNPs by the four GWAS packages. Also, the linkage disequilibrium analyzed confirmed this replicability. Furthermore, when the Naive GWAS model was used to analyze the same dataset, MultiGWAS showed all four tools simultaneously detected the SNP c1_8019 as a significant associated SNP, highlighting it as a reliable association. (see Supplemental Material S1 and S2 at https://github.com/agrosavia-bioinfo/multiGWAS/tree/master/docs.)

Two pairs of SNPs resulted in LD, c2_8019 with c2_25471 and c2_45606 with c2_45611, named by MultiGWAS as LD_SNP1 and LD_SNP2, respectively (table of Figure 7.a). The Venn diagram (Figure 7.b) shows that almost one SNP of the pairwise SNPs in LD was detected by the four GWAS pack-ages, showing the replicability of the SNPs in the four packages. Moreover, the chord diagrams show that most of the best-ranked SNPs were in chromosome 10 (Figure 7.c). Finally, the best-ranked SNP’s heatmaps show visible differences that related the association of the genotype with the phenotype for tuber shape (Figure 7.d).

The Manhattan plot for each GWAS package showed that four packages found that the associated SNP location (i.e., SNP above the blue line) was chromosome 10 (Figure 8). From them, are significant by GWASpoly and SHE-sis (SNPs above the red line). Both SNP groups, the strong associated and the significant, are present in both the shared table and Venn diagram (Figure 6).

**Figure 8:**
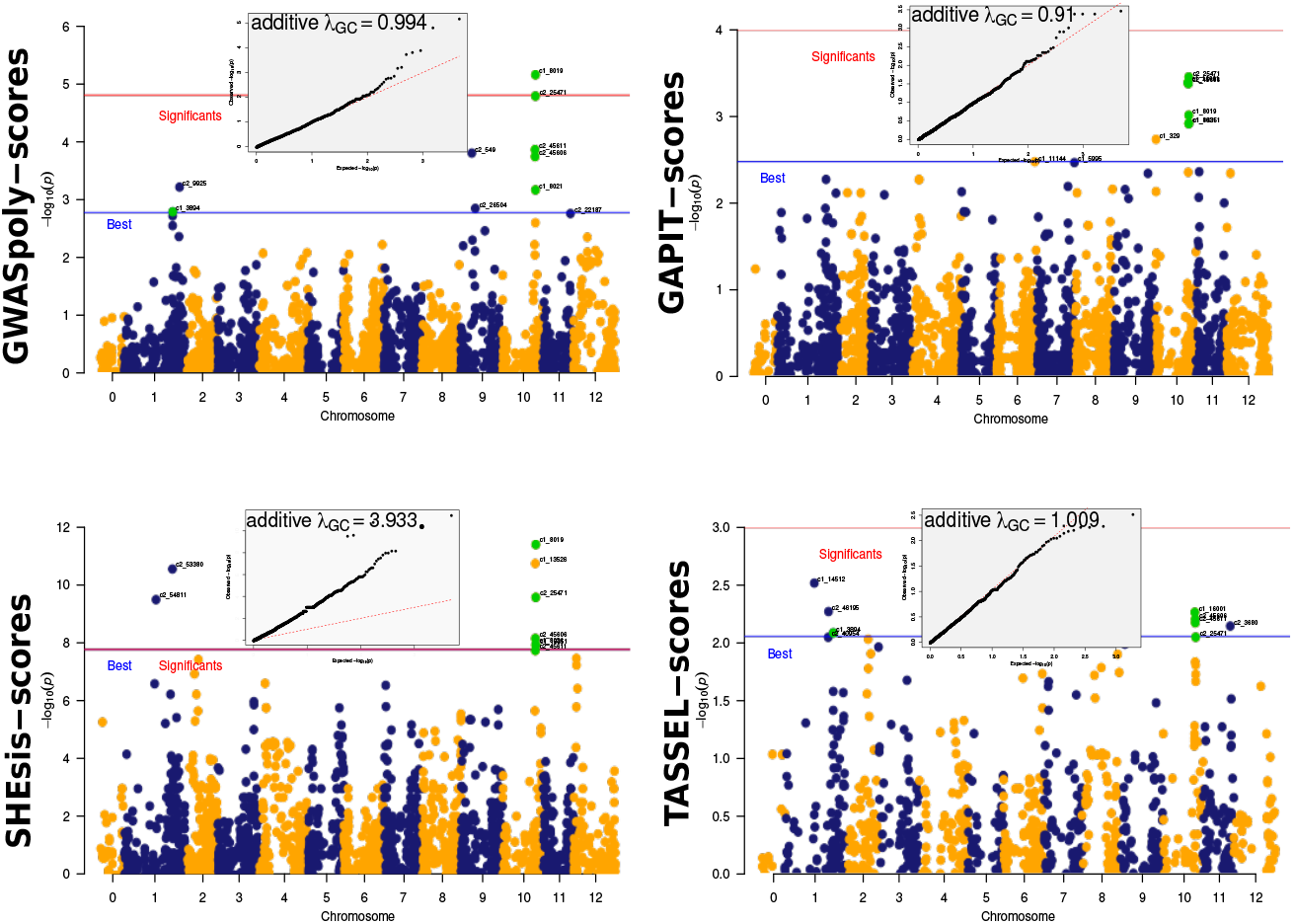
Associations for each GWAS package. MultiGWAS shows the associations identified by the four GWAS packages using Manhattan and QQ plots. The tetraploid potato data showed several SNPs shared between the four GWAS packages (green dots). The best-ranked SNPs are above the blue line, but only GWASpoly and SHEsis identified significant associations (SNPs above the red line) for this dataset. However, the inflation factor given by SHEsis is too high (λ = 3.9, at the top of the QQ plot), which is observed by the high number of SNPs deviating from the red diagonal of the QQ plot.

Additionally, for most GWAS packages, except for SHEsis, the majority of observed *p-values* corresponded to the expected *p-values*, as it is shown in the QQ plots generated from the associations found for each package (QQ plots above Manhattan plots in Figure 8). For SHEsis, its genomic inflation factor λ was far above 1.0, meaning that its calculated scores were inflated, and explaining because SHEsis does not control for population structure and relatedness.

### 5.2 MultiGWAS performance in simulated data

For MultiGWAS, we present the results using different sets to evidence the effect of replicability in the performance (**MultiGWAS_1**: predicted by one software, **MultiGWAS_2**: predicted by two software, **MultiGWAS_3**: predicted by three software, and **MultiGWAS_4**: predicted by four software).

For the additive effect simulation, GWASpoly (green) and SHEsis (blue) had the best performance based on True Positive Rate (TPR) and True Negative Rate (TNR) in the detection of the best-ranked SNP. The two diploid software GAPIT and TASSEL, have similar results but lower performance in both statistics. In parallel, MultiGWAS performance change depending on the number of software involved in the predicted SNP intersection; the TPR was progressively lower, and TNR progressively higher. Consequently, in the two more restrictive cases (i.e., the intersection of predicted SNP by three and all the four software, MultiGWAS_3 and MultiGWAS_4 respectively),the TPR was similar to obtained by TASSEL and GAPIT. However, the TNR was higher than even GWASpoly and SHEsis (Figure 9.a)

**Figure 9:**
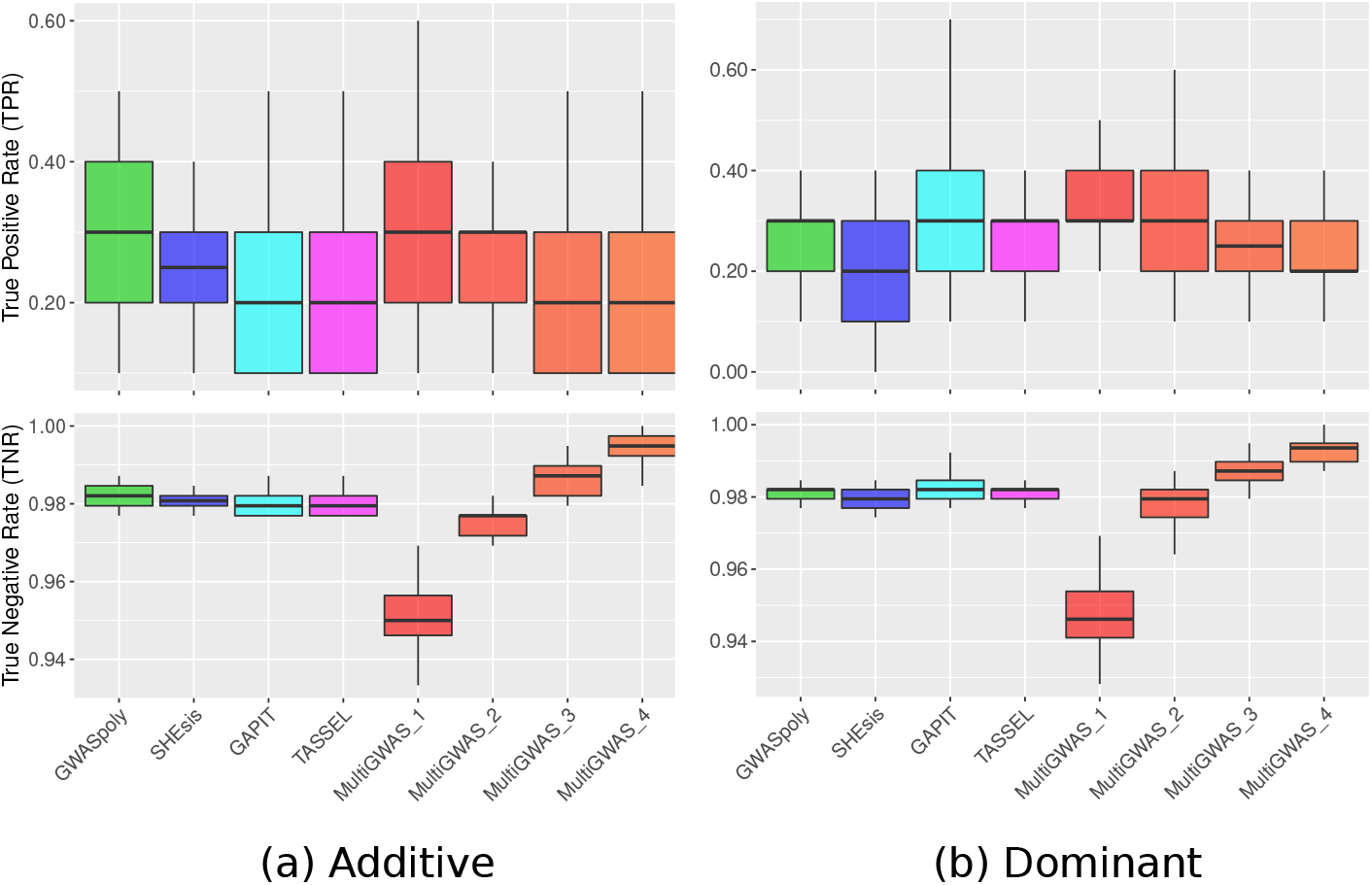
Boxplots for the true-positive rates (TPR) and true-negative rates (TNR) for the top SNPs (i.e., the N best ranked) identified for each GWAS software using simulated datasets either under (a) additive or (b) dominant inheritance model after 50 replicates. Each panel compares MultiGWAS with each of the four software that integrate it (i.e., GWASPoly, SHEsis, GAPIT, and TASSEL). The MultiGWAS results are separated into four groups thus: MultiGWAS_1, predicted by one tool. MultiGWAS_2, predicted by two tools. MultiGWAS_3: predicted by three tools, and MultiGWAS_4: predicted by four tools.

For the dominant effect simulation, GAPIT (cyan) tends to have a higher TPR than the other three software. Moreover, SHEsis had a lower value of both TPR and TNR since it was designed only to detect associations with additive effects. Comparing these four software’s performance with a wrapping tool as MultiGWAs, it had a similar performance to the additive effect simulation. As the more restrictive the intersection is, the TPR was progressively lower and TNR progressively higher. However, in the two more restrictive cases (i.e., the intersection of predicted SNP by three and all the four software, MultiGWAS_3 and MultiGWAS_4 respectively), the TNR was higher than all the four software (Figure 9.b)

In the case of significant SNPs, for additive effects, SHEsis (blue) has the highest TPR but has the lowest TNR, suggesting that SHEsis probably is over-estimating the significative *p-values* association. Therefore true and false associations are reaching the significance threshold. In comparison, MultiGWAS_4, GWASpoly, and GAPIT are more conservative, with closer TPR but high TNR (Figure 10.a).

**Figure 10:**
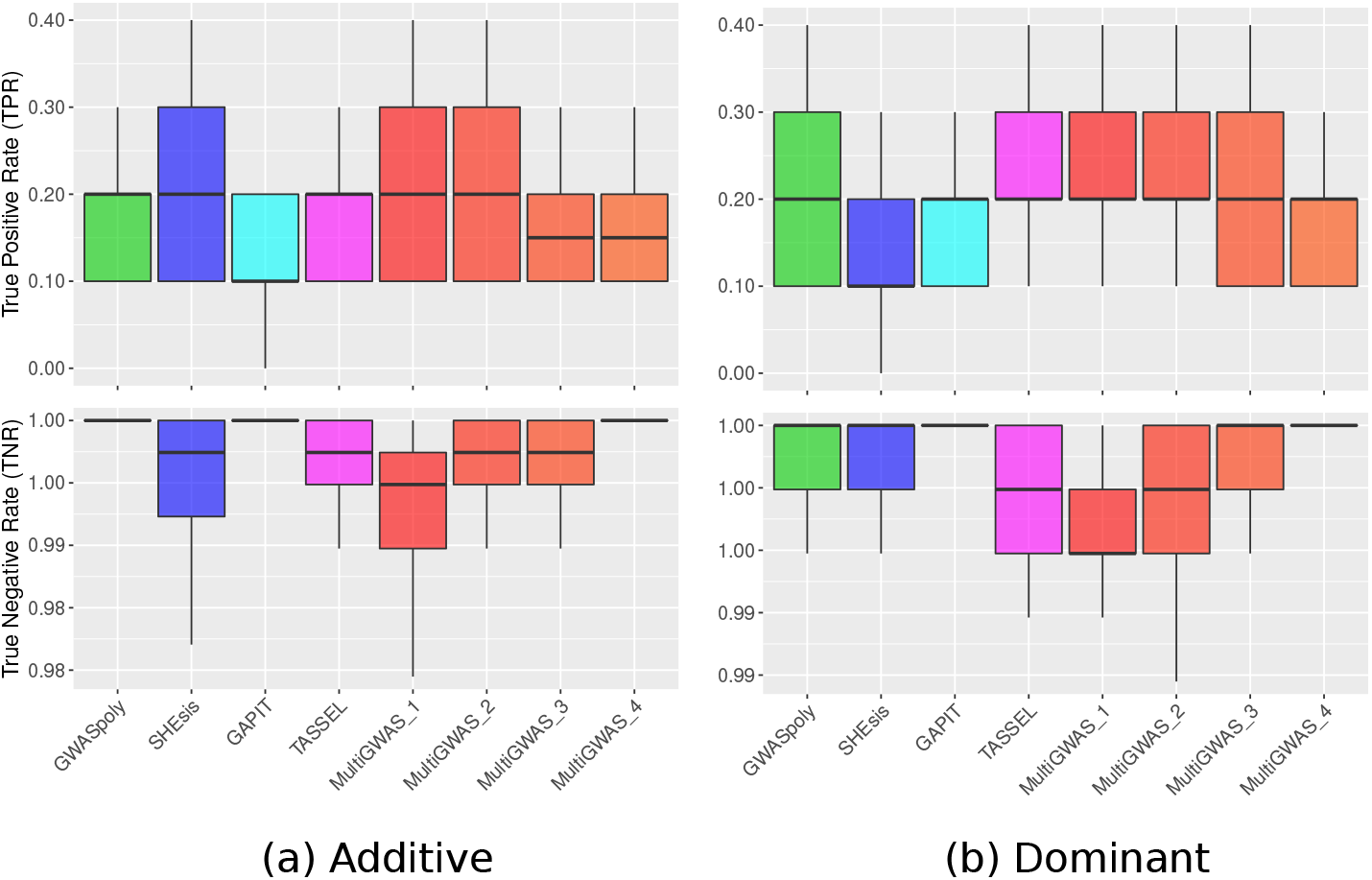
Boxplots for the true-positive rates (TPR) and true-negative rates (TNR) for the identification of significant SNP identified for each GWAS software using simulated datasets either under (a) additive or (b) dominant inheritance model after 50 replicates. Each panel compares MultiGWAS with each of the four software that integrate it (i.e., GWASPoly, SHEsis, GAPIT, and TASSEL). The MultiGWAS results are separated into four groups thus: MultiGWAS_1, predicted by one tool. MultiGWAS_2, predicted by two tools. MultiGWAS_3: predicted by three tools, and MultiGWAS_4: predicted by four tools.

For dominant effects simulation, GWASpoly had the higher TPR with a lower TNR. Thus, this software was the most sensitive detecting significative associations, but at the same time, it was one of the least specific. In comparison, MultiGWAS_4 and GAPIT had TPR slightly lower than of GWASpoly, but with the highest TNR. This pattern suggests that both are less sensitive in detecting significative association but more specific than GWASpoly. Therefore, MultiGWAS_4 provides very accurate associations (Figure 10.b).

## 6 Discussion

The reanalysis of both potato and simulated data with MultiGWAS showed that this wrapping tool is handy to improve the GWAS in both diploid and tetraploid species for additive and dominant gene action effects. Through MultiGWAS performance, we could test its effectiveness to answer some of the challenges of analyzing polyploid organisms. They include integrating and replicating among parameters and software, the diploidization of polyploid data, and the incorporation of different inheritance mechanisms [Dufresne et al., 2014].

The main advantage of MultiGWAS is that it replicates the GWAS analysis among four software and integrates the results obtained across software, models, and parameters. Therefore, in MultiGWAS, users do not have to choose between specificity or sensitivity because they can observe their effect in the analysis within the same wrapping environment.

Another difficulty for replication among software is the variability of formats for the genomic input data. MultiGWAS receives the genotype data in five different formats, including two software outputs used to call polyploid allele dosage. Currently, the most common format for next-generation sequencing variant data is the VCF (Variant Call Format) [Danecek et al., 2011][Ebbert et al., 2014]. One of the advantages of VCF is its versatility in summarizing important genome information for hundreds or thousands of individuals and SNP, including information about ploidy levels. MultiGWAS simplifies using the GWAS software because it allows the VCF files as an input (but see VarStats tool in VTC).

Moreover, the MultiGWAS is the unique wrapping tool we are aware of that facilitates understanding the effect of diploidizing the tetraploid data in the analysis’s performance directly. The SNP profile allows identifying what the significant associations detected by more than one software are. Furthermore, although MultiGWAS checks for significative SNPs based on the *p-value*, it is essential to go back to the data and check if the SNP is a true association between the genotype and phenotype. For this purpose, the SNP profile gives visual feedback for the accuracy of the association.

Furthermore, the MultiGWAS allows comparing among the gene action models that offer GWASPoly and TASSEL. GWASpoly [Rosyara et al., 2016] provides models of polyploid gene models action, including additive, diploidized additive, duplex dominant, simplex, and general. On the other hand, TASSEL [Bradbury et al., 2007] also models different gene action types for general, additive, and dominant diploids. To choose among models, we propose an automatic selection of the gene action model for both tools based on a balance between three criteria: the inflation factor, the replicability of identified SNPs, and the significance of identified SNPs. This inflation index is a new tool for comparison that does not offer either GWASPoly or TASSEL. This automatic strategy will help to understand the gene action model for the trait of interest. Although the main focus is on the resultant SNPs, the model has assumptions that reflect a specific phenotype’s gene actions.

Finally, MultiGWAS, through the active comparison among models, addresses the search of the inheritance mechanisms by comparing among two designed for polysomic inheritance software [Rosyara et al., 2016, Shen et al., 2016] with two software for disomic inheritance [Purcell et al., 2007, Bradbury et al., 2007]. Understanding the inheritance mechanisms for polyploid organisms is an open question. For autopolyploids, most loci have a polysomic heritage. However, sections of the genome that did not duplicate lead to disomic inheritance for some loci [Ohno, 1970, Lynch and Conery, 2000, Dufresne et al., 2014]. Thus it is a valuable tool for researchers because it looks for significant associations that involve both types of inheritance.

### 6.1 Future remarks

The evolution and population genomics of polyploids is an exciting novel area of research. The advancement of next-gen sequencing techniques produces more empirical polyploid data in different model and non-model organisms [Ekblom and Galindo, 2011, Ellegren, 2014].

Many assumptions developed for diploids in the GWAS analysis do not apply entirely for polyploids [Dufresne et al., 2014]. Fortunately, in the last five years, different models to calculate several parameters for population genomics on polyploids are testing and developing in simulated and empirical data [Meirmans et al., 2018, Hardy, 2016, Blischak et al., 2016].

For MutiGWAS, we started with the most simple ploidy, such as tetraploids. Nevertheless, future MultiGWAS versions should include more complex ploidies than tetraploids and the explicit calculation of parameters either for filtering polyploid data before GWAS analysis or complementing other populations genomics’ parameters of the data analyzed.

## 7 Acknowledgements

This research was possible thanks to AGROSAVIA five-years macroproject entitled *Investigación en conservación, caracterización y uso de los recursos genéticos vegetales*.

We thanks to the Minister of Science, Technology and Innovation of the republic of Colombia (previously COLCIENCIAS), for supporting the postdoc-toral researcher L. Garreta at AGROSAVIA during 2019-2020 under the supervision of ICS and PHRH (Grant number 811-2019). The editorial of AGROSAVIA gave for finatial supporting for this publication. Finally to Andres J. Cortés for his valuable comments to improve this manuscript.

## 8 Author Contributions

LG, ICS, and PHRH conceived the idea. LG developed MultiGWAS. MP tested MultiGWAS. All authors wrote and approved the final version of the manuscript.

